# The type and source of reactive oxygen species influences the outcome of oxidative stress in cultured cells

**DOI:** 10.1101/2021.02.16.431512

**Authors:** Steffi Goffart, Petra Tikkanen, Craig Michell, Trevor Wilson, Jaakko L.O. Pohjoismäki

**Author notes:** Equal contribution (Steffi Goffart, Petra Tikkanen).

## Abstract

Oxidative stress can be modeled using various different experimental approaches, such as exposing the cells or organisms to oxidative chemicals. However, the actual effects of these chemicals, outside of the immediate measured effect, have attracted relatively little attention. We show here that three commonly used oxidants, menadione, potassium bromate and hydrogen peroxide, while known to function differently, also elicit different types of responses in cultured cells. While cells response to menadione and bromate exposure mainly by an integrated stress response, hydrogen peroxide has more indirect effects. Primary oxidative stress does not induce DNA repair or antioxidant defense mechanisms. However, cells with previous experience of oxidative stress show adaptive changes when the stress is renewed. Our results urge caution when comparing studies using different sources of oxidative stress or generalizing the findings of these studies to different tissue or oxidant types.

## Introduction

Electron-deficient atoms or molecules, having unpaired valence electrons, can draw electrons from others, making them chemically highly reactive (1). Consequently, these species are commonly known as free radicals, causing significant harm to biological macromolecules such as proteins, lipids and nucleic acids (2). Because of the abundance and chemical nature of oxygen, reactive oxygen species (ROS) are the most important naturally occurring type of free radicals. In cells, the main source of ROS is the mitochondrial electron transport chain, where uncontrolled leakage of electrons through complexes I and III can generate superoxide (O_2_·^-^) from oxygen (3). Superoxide is then either spontaneously dismutated or actively converted to hydrogen peroxide (H_2_O_2_) by the mitochondrial superoxide dismutase (SOD2) and further to water (H_2_O) by catalase or glutathione peroxidases (GSHs). Although O_2_·^-^ and H_2_O_2_ are not especially reactive themselves, their reactions with transition metals generate extremely reactive hydroxyl radicals (OH·), which cause the majority of oxidative damage in cells (4).

The effects of ROS on cells have been studied intensively because of their importance in human pathologies, cellular signaling and ageing (2,3,5). While oxidative stress can be experimentally increased by genetic manipulation of antioxidant defenses (6) or ionizing radiation (7), the most common method is to expose cells or animals to chemicals capable of generating ROS. These compounds can be oxidants themselves, block the mitochondrial electron transport chain (ETC) or bypass respiratory complexes by transferring electrons directly to oxygen. An example of the latter is menadione, a quinone and vitamin K analogue, which can transfer electrons from the ETC complex I directly to oxygen, generating superoxide (8). Menadione has been used for a long time in a broad spectrum of studies to induce oxidative stress, cell damage and death (9–12). Similarly, H_2_O_2_ has offered an easy experimental source of ROS stress and is widely used in experimentation (13). H_2_O_2_ is notoriously difficult to handle reproducibly, has relatively short half-life and is prone to both enzymatic as well as chemical elimination in medium or in cells (13, 14). Generally, the H_2_O_2_ quantities required to induce oxidative damage are an order of magnitude higher than for other ROS stressors.

Our group has been interested in the consequences of oxidative damage on mtDNA, which we have modeled mainly by exposing cells to potassium bromate (KBrO_3_) (15–17). In contrast to menadione, bromate can directly oxidize macromolecules, including DNA in cells (16, 18). In mitochondria, potassium bromate treatment increases the levels of 8-oxoguanine (8-oxoG) modifications on mtDNA (16) and causes persistent changes in its replication mechanisms (15, 16). Interestingly, menadione and H_2_O_2_ treatments have different outcomes on mtDNA in our hands. However, it can be argued that menadione-induced oxidative stress is physiologically more relevant as it is caused by superoxide originating from the electron transport chain, recapitulating the natural sources during intense oxidative metabolism (3).

In order to understand the differences between oxidative stressors and evaluate their relevance for modelling various physiological sources of ROS damage, we have compared the effects of menadione, potassium bromate and H_2_O_2_ on cellular responses. While menadione elicits intrinsic ROS by producing O_2_·^-^ within the mitochondria, the effects of KBrO_3_ and H_2_O_2_ can cause direct oxidative damage at the cell surface or upon entry into the cells. In contrast to bromate, superoxide and H_2_O_2_ are not necessarily functioning as oxidizing agents themselves, but might require conversion to hydroxyl radicals. As the superoxide resulting from menadione treatment can be converted to H_2_O_2_, one would expect the two oxidants to have comparable effects on cells. Interrogating oxidative stress signaling pathways, DNA damage responses and changes in the gene expression patterns, we note that all three compounds cause rather dissimilar effects on cells. Our study emphasizes the notion that not all ROS stressors are equal, and that caution should be exercised when generalizing the findings.

## Results

In order to get a first impression on how menadione, KBrO_3_ and H_2_O_2_ might differ in their action as ROS stressors, we compared the ability of these compounds to oxidize mitochondrial and cytoplasmic redox-sensitive marker molecules. Interestingly, high concentrations of H_2_O_2_ had little effect on cytoplasmic or mitochondrial ROS, whereas menadione-induced roGFP oxidation in both compartments (Figure 1A). Somewhat unexpectedly, KBrO_3_-induced roGFP oxidation was specific for the mitochondrial compartment. A commonly used marker of mitochondrial oxidative stress, MitoSOX, reacted only with menadione (Figure 1B). This is not surprising as MitoSOX is relatively specific for superoxide (19), which is generated by menadione but not by the other two chemicals.

**Figure 1.**
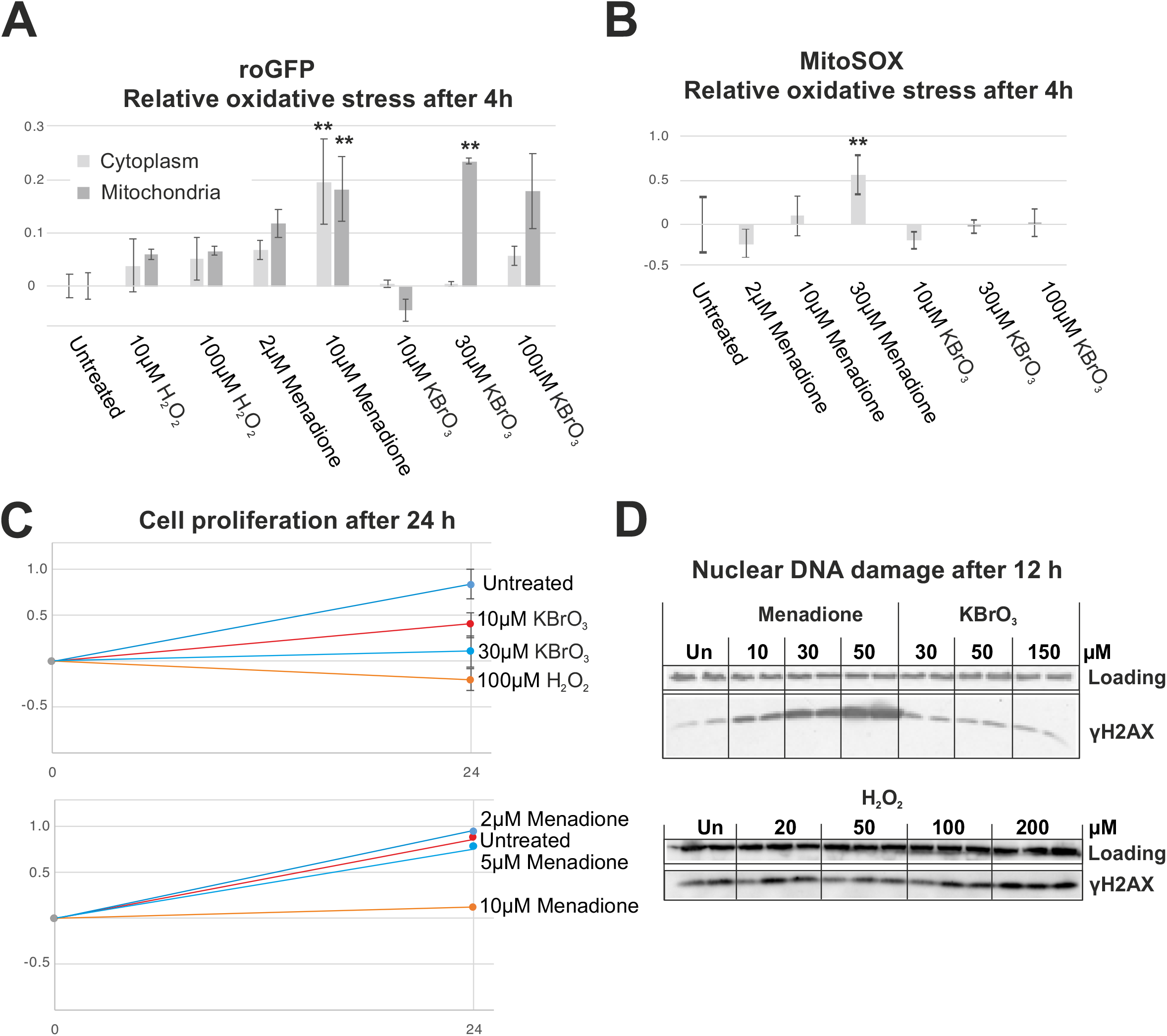
Oxidative stress, nuclear DNA damage and cell proliferation after menadione, KBrO_3_ and H_2_O_2_ exposure. (A) Cytoplasmic and mitochondrial oxidative stress in HEK293T cells measured as a relative change in the roGFP excitation spectrum after 4 h treatment with different oxidant concentrations. (B) Mitochondrial superoxide levels in HEK293T cells after 4 h treatment with different oxidant concentrations. (C) Cell proliferation during 24 h exposure to different oxidant concentrations. (D) Activation of nuclear DNA damage signaling – phosphorylation of histone H2AX (gH2AX) – in HEK293T cells after 12 h treatment with different oxidant concentrations. Vinculin was used as loading control for the Western blots.

Despite the differences in the observed oxidative stress, the highest concentrations of all drugs were able to stop cell proliferation (Figure 1C). As of note, menadione concentrations above 50 μM and H_2_O_2_ above 200 mM concentrations killed the HEK293T cells effectively, whereas KBrO_3_ was well tolerated at the observed time points. While menadione-induced nuclear DNA double-strand break signaling was activated already at relatively low concentrations (Figure 1D), this was not observed even with highest concentrations of KBrO_3_ and H_2_O_2_.

Next, we focused on the activation of integrated stress response (ISR) upon ROS stress. There are a number of different entries into ISR, but its key downstream signaling protein is ATF4, whose translation is increased upon ISR activation (20). While ATF4 was substantially upregulated after 12 h of menadione exposure, KBrO_3_ and H_2_O_2_ showed again no or little effect (Figure 2A). However, when we studied the timing of ISR activation in more detail, we noted that KBrO_3_ caused an increase in ATF4 aftar more than 12 h, while H_2_O_2_ caused only a transient increase in ATF4 levels at 4 h (Figure 2B).

**Figure 2.**
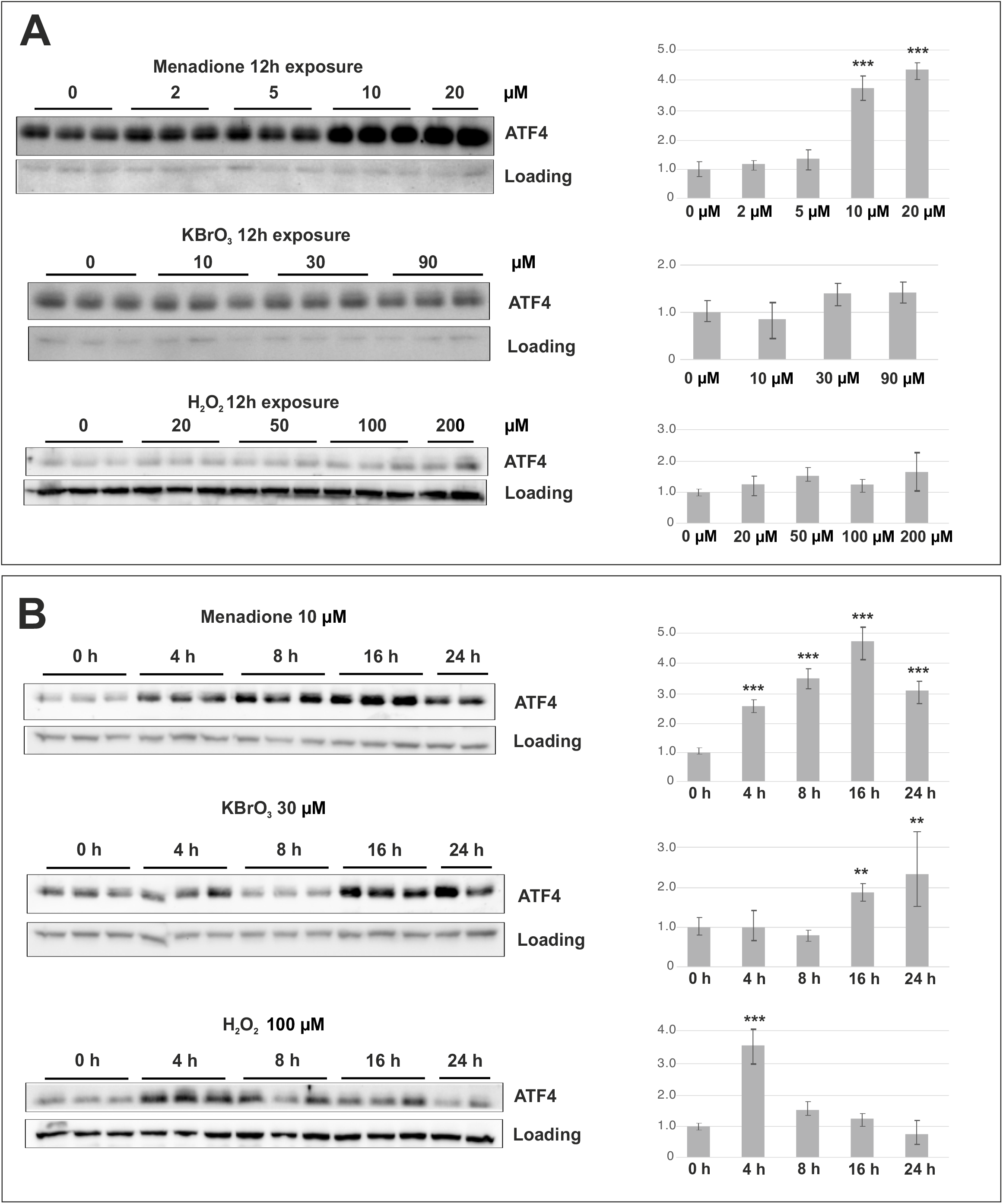
Integrated stress response (ISR) in HEK293T cells exposed to menadione, KBrO_3_ or H_2_O_2_. (A) ATF4 levels after 12 h exposure to increasing concentrations of oxidants. Note the dramatic increase after Menadione treatment. (B) Menadione causes a rapid and persistent induction of ISR, whereas KBrO_3_ treated cells show a delayed response only after 16 h exposure. The H_2_O_2_ treatment results in a transient increase in ATF4 levels after 4 h exposure. VDAC and actin were used as a loading control. * *p*<0.05, ** *p*<0.01, *** *p*<0.001 ANOVA/Tukey, compared with the untreated control.

Despite their differential effects on DNA damage response (Figure 1D) and ISR activation (Figure 2), 10 μM menadione, 30 μM potassium bromate, and 100 mM hydrogen peroxide were all able to halt cell proliferation at 24 h (Figure 1C). To obtain a more global view on the effects of these chemicals and cause of the growth arrest, we performed a transcriptome analysis using an RNA sequencing approach. The 24 h timepoint was chosen as a treatment end-point result to compare differences and similarities from the various oxidants. As potassium bromate is not rapidly turned over like H_2_O_2_ and is also not known to interfere with the mitochondrial respiratory chain as menadione, thus avoiding downstream effects on the overall cellular metabolism, we took it as an example oxidant to study the recovery and subsequent response to a repeated ROS insult as an additional experiment. As an additional aspect, we have shown that KBrO_3_ induces specific changes in mtDNA replication, which we have not observed with the other two oxidants.

A 24 h treatment with menadione and potassium bromate resulted in a strikingly similar effect on gene expression, whereas H_2_O_2_ treatment caused a markedly different reaction in cells (Figure 3A). Cells treated with KBrO_3_ and let to recover for additional 24h were comparable to untreated cells, demonstrating that the oxidative exposure did not result in permanent alteration of gene expression. Interestingly, the gene expression changes in cells retreated with potassium bromate after the initial recovery did not correspond to cells treated only once. The similarity of the first time KBrO_3_ and menadione treatment was caused by substantial reduction in general transcription activity (Figure 3B, C), whereas the hydrogen peroxide treatment resulted in differential up- and downregulation of dozens of genes (Figure 3D). Genes significantly upregulated in hydrogen peroxide-treated cells included antioxidant defense enzymes, such as *SOD1* as well as glutathione peroxidases *GPX1* and *GPX8.* Interestingly, neither mitochondrial *SOD2* nor catalase *(CAT),* responsible for H_2_O_2_ elimination in mitochondria, were affected. 10 μM menadione caused a slightly stronger reduction in transcript levels than 30 μM potassium bromate (Figure 3E).

**Figure 3.**
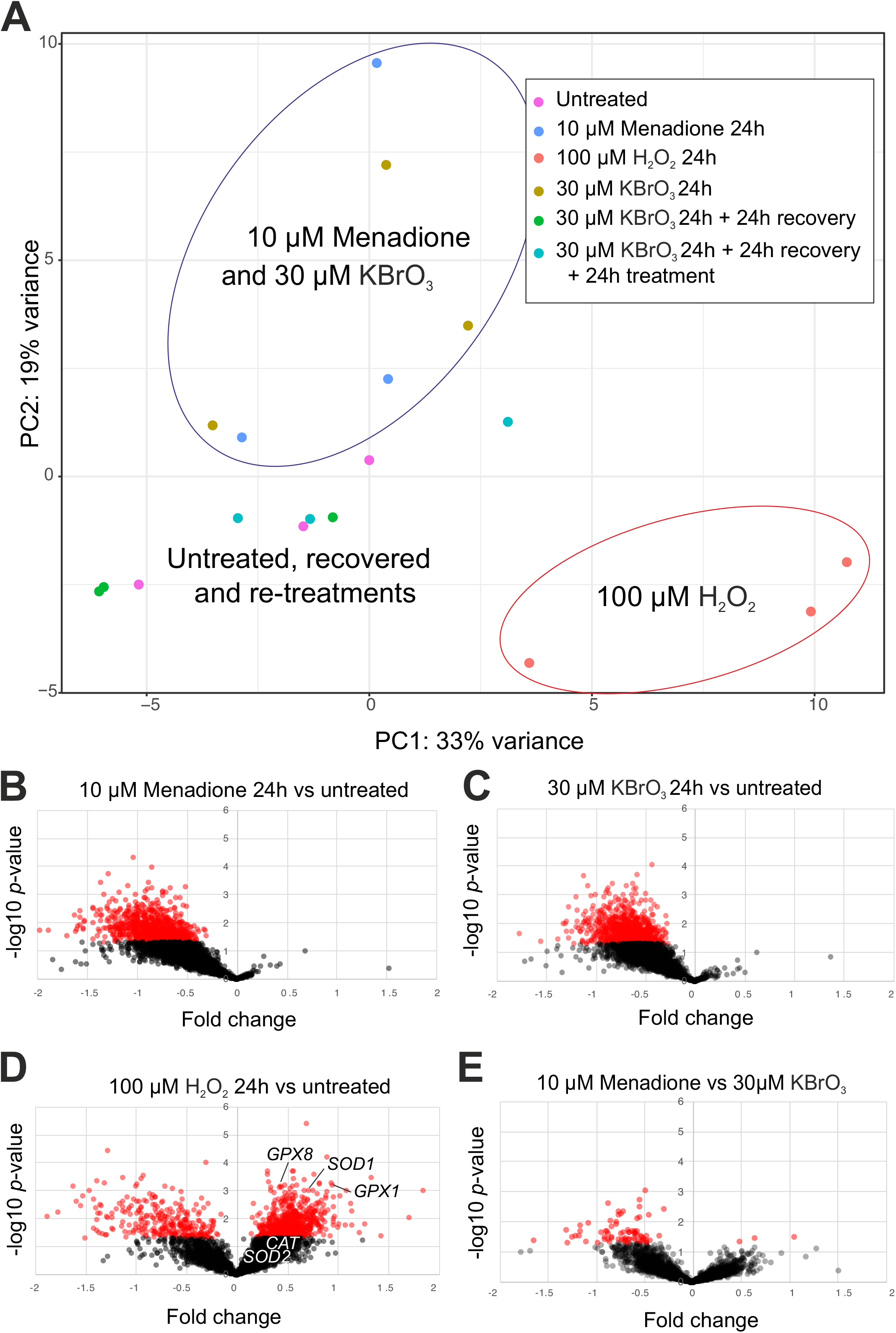
Global gene expression changes during and after oxidative stress. (A) Principal component analysis (PCA) of the treated cells as indicated in the legend. Menadione and KBrO_3_ exposure cause similar changes in the transcriptome, whereas H_2_O_2_ treated cells differ from these as well as control cells. Untreated controls, cells recovered from KBrO_3_ exposure for 24h and cells re-treated with KBrO_3_ after recovery also cluster together. (B) A volcano plot showing down-(left) and up- (right) regulated genes after 24 h menadione treatment. The x-axis indicates the inverse log10 t-test *p*-value with *p* < 0.05 indicated in red. (C) The KBrO_3_-treated cells show a similar, general reduction in gene expression as the menadione treated cells. (D) In contrast, H_2_O_2_ treatment results in differential up- and downregulation of dozens of genes. Antioxidant defense genes, such as *SOD1* and *GPX*s are among the upregulated genes. (E) 10 μM menadione treatment causes a more exaggerated reduction in transcriptional activity than 30 μM KBrO_3_.

Despite the dramatic differences in the impact on the global gene expression of menadione and KBrO_3_ compared to the H_2_O_2_ treatment, the effects of the three different oxidant exposures had some common outcomes on negatively regulated genes (Figure 4A). Notably, not a single positively regulated gene was shared between the treatments. All three treatments resulted in downregulation of cell cycle (Figure 4B), as evident also from the cessation of cell proliferation (Figure 1C). Both menadione and potassium bromate treatments also resulted in general downregulation of gene expression (Figure 4C), which was not evident after hydrogen peroxide treatment. In addition, KBrO_3_-treated cells showed specific impacts on translation, specifically the downregulation of translation initiation, nonsense-mediated decay and post-translational modification (Figure 4D), which were not seen in cells treated with menadione. In fact, apart for general transcription inhibition, menadione treated cells showed only a weak impact on other specific cellular processes (Figure 4E). In contrast to menadione and KBrO_3_, H_2_O_2_-treated showed specific and substantial upregulation of a number of cellular processes, including amino acid metabolism, protein catabolism, hypoxia response and mitochondrial biogenesis (Figure 4F).

**Figure 4.**
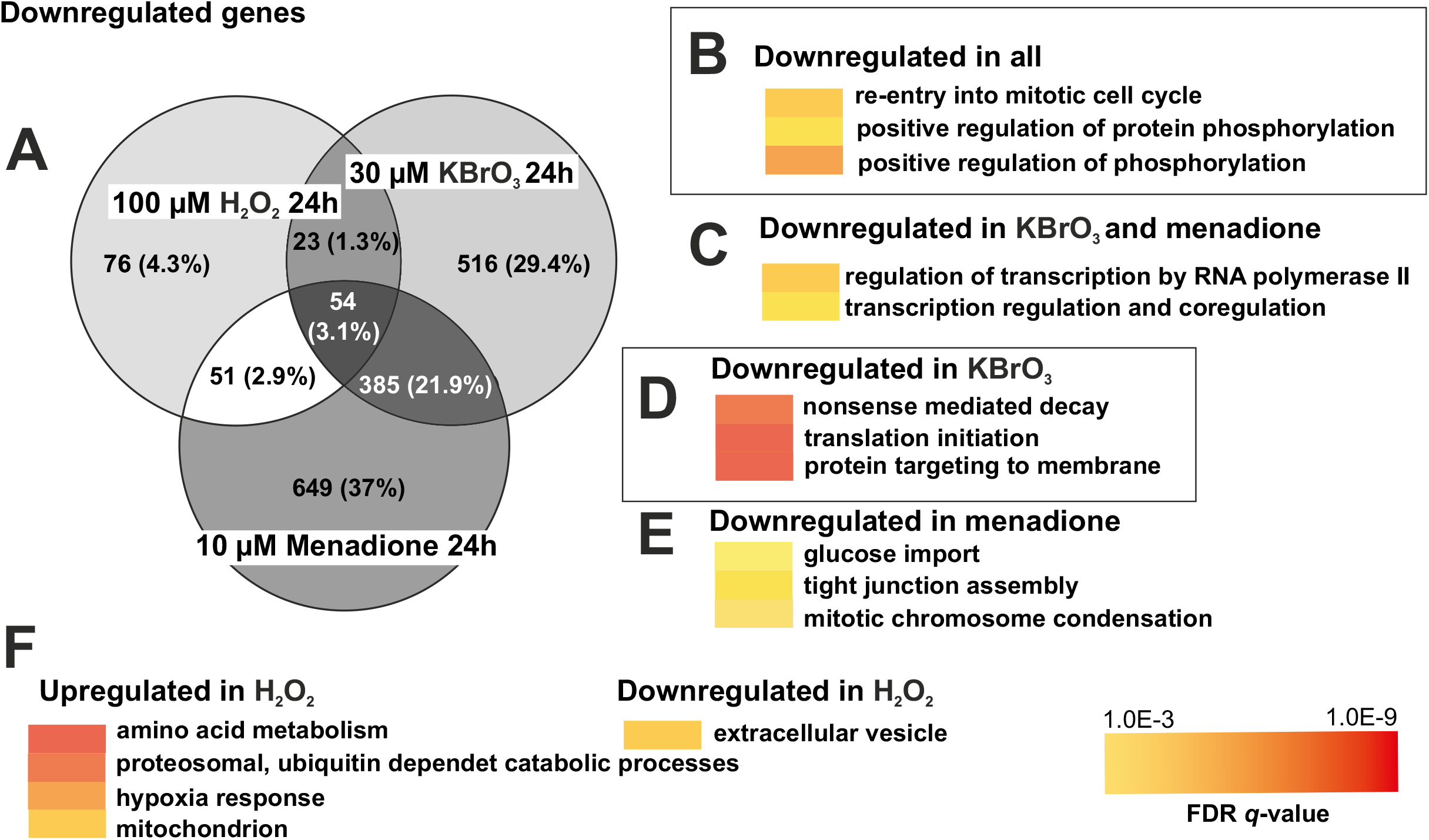
Different oxidative insults have common features. (A) A Venn diagram showing the number of common and private, significantly down-regulated genes after menadione, KBrO_3_ or H_2_O_2_ treatment. (B) Among the 54 genes downregulated after the exposure to each of the three chemicals were genes that are involved in cell cycle regulation and cell signaling. (C) Both menadione- and KBrO_3_-treated cells showed general inhibition of transcription, while (D) KBrO_3_ alone induced also downregulation of translation. (E) Menadione had relatively few privately affected processes, among these glucose import. The 76 genes downregulated in H_2_O_2_-treated cells did not show any significant enrichment of specific cellular processes. (F) In contrast, H_2_O_2_-treated cells showed a significant upregulation of genes involved in amino acid metabolism, hypoxia response and protein catabolism.

As pointed out earlier, ATF4-mediated integrated stress response can be triggered by a number of upstream signals. The most logic one upon oxidative stress would be the unfolded protein response (21). Oxidation of disulfide bonds in proteins results in the accumulation of misfolded proteins, triggering PERK, a transmembrane protein kinase that phosphorylates the α-subunit of translation initiation factor 2 (eIF2α). Phosphorylation of eIF2α in turn increases the translation of *ATF4* mRNA while otherwise generally inhibiting protein synthesis. Interestingly, Menadione exposure did not increase eIF2α phosphorylation (Figure 5A) in HEK293T cells in similar fashion as UV exposure (Figure 5B). However, menadione treatment was able to overcome the decrease in the eIF2α phosphorylation caused by PERK inhibitor, maintaining steady-state levels of the phosphorylated protein (Figure 5A). Oddly, the same was not observed for UV treatment (Figure 5B).

**Figure 5.**
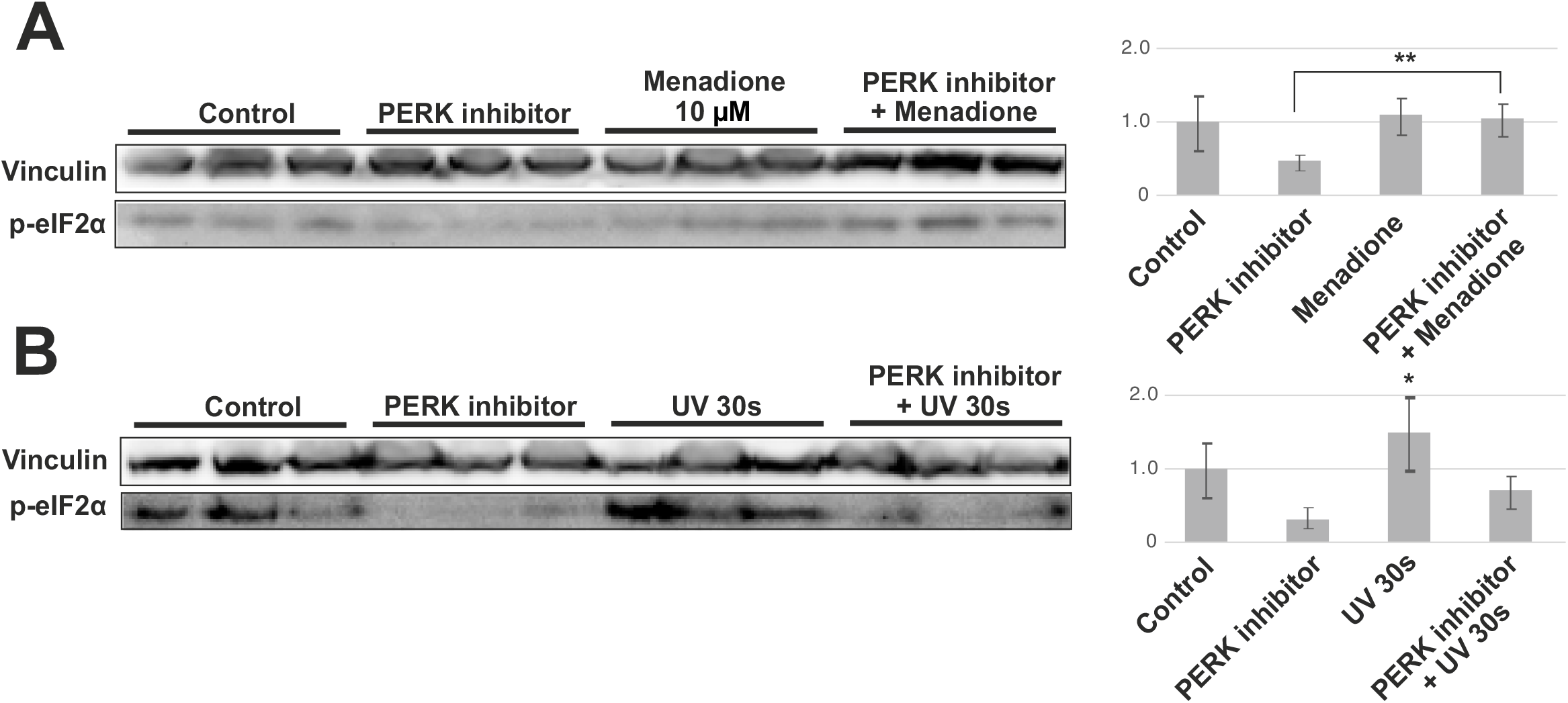
ISR upstream signals after menadione treatment. (A) Despite rapid upregulation of ATF4 levels, eIF2α phosphorylation is not increased in cells treated with menadione for 8 h. While a PERK inhibitor abolishes eIF2α phosphorylation in untreated cells, steady-state levels are maintained in menadione-treated cells also in the presence of the inhibitor. (B) To validate the functionality of the antibody as well as the existence of PERK-eIF2α signaling in HEK293T cells, the cells were exposed to 1.34 mJ/cm^2^ UVB (305 nm) for 30 s. Addition of PERK inhibitor 4 h prior the exposure prevented eIF2α phosphorylation. Vinculin was used as a loading control. * *p*<0.05, ** *p*<0.01 ANOVA/Tukey.

As we have used potassium bromate successfully to induce mtDNA damage and subsequent changes in the replication mechanisms, we next compared the effects of the different oxidants on the replication intermediates 3 kb downstream of the main replication origin (Figure 6). The reason to investigate this region is that it allows to compare the partly single-stranded mtDNA replication intermediates arising from the house-keeping, strand-asynchronous replication mechanism with double-stranded DNA replication intermediates, that are typical for genomic stress (15–17). In line with our previous work, only the KBrO_3_ treatment was able to elicit a change in the mtDNA replication pattern after 24 h treatment.

**Figure 6.**
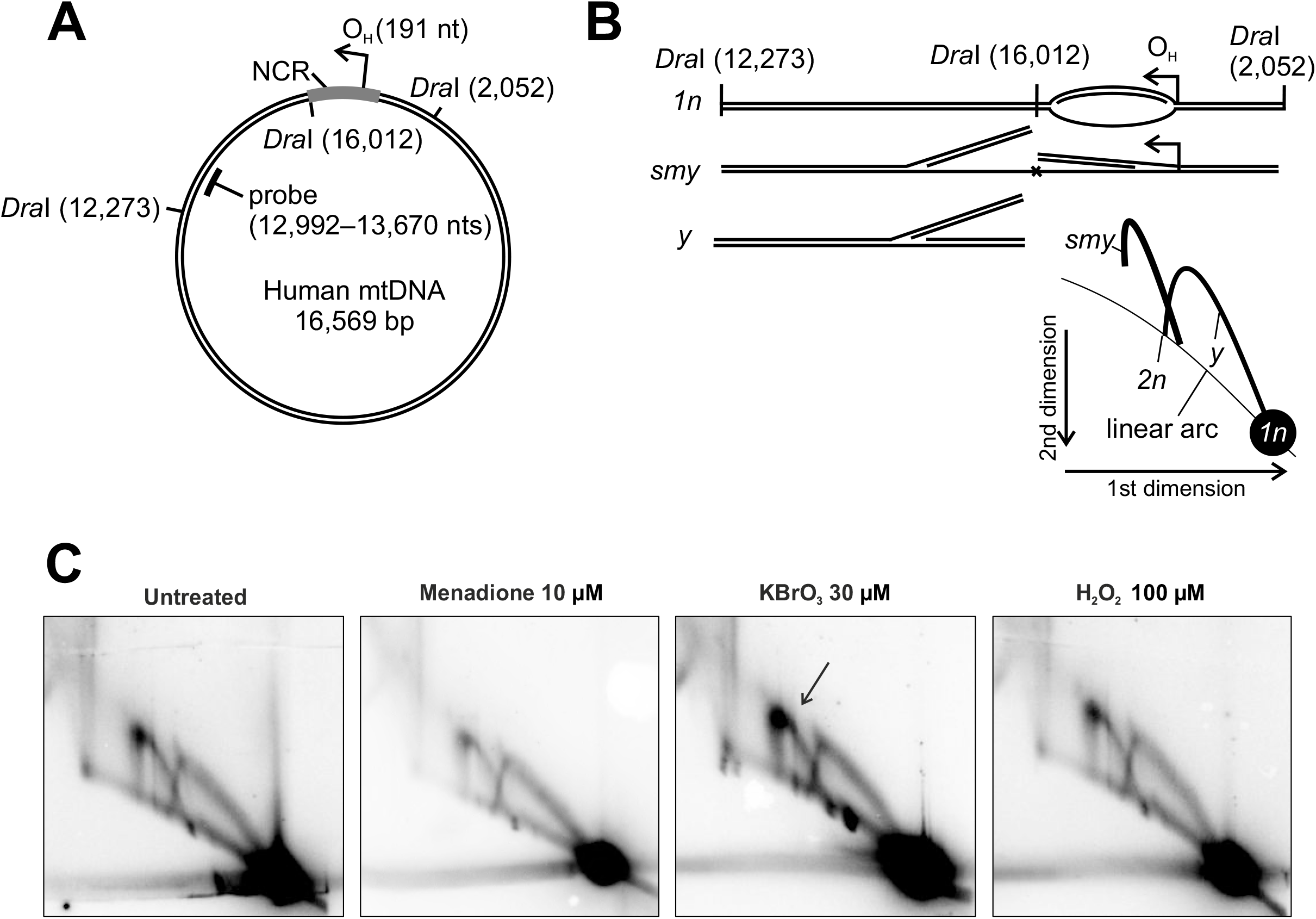
Modulation of mitochondrial DNA replication during oxidant exposure. (A) Schematic diagram of human mtDNA showing the investigated genomic region. (B) Interpretation of the 2D-AGE pattern. The probed fragment is downstream of the main replication origin OH. Restriction site blockage due to single-strandedness or RNA incorporation on the lagging strand will give rise to slow moving y-arc (*smy*) whereas doublestranded DNA replication intermediates migrate the the y-arc (*y*). (C) 2D-AGE of *Dra*I digested mtDNA from untreated control cells, cells treated with 10 μM menadione, cells treated with 30 μM KBrO_3_ and from cells treated with 100 μM H_2_O_2_ for 24h. Arrowhead points to the increase in the partly single-stranded mtDNA replication intermediates in KBrO_3_-treated cells.

We were also interested to see if an initial exposure to oxidative stress is able to ameliorate the effects of a second ROS insult, a phenomenon known as (mitochondrial) hormesis (3, 22). Based on our transcriptome analyses, 293 cells recovered well within 24 h after an initial insult with 30 μM potassium bromate (Figure 3A, Figure 7A). Some genes remained downregulated compared to untreated cells, but no significant regulation of specific cellular processes was detected in the GO-term analyses. Interestingly, re-treatment of the recovered cells with the same concentration of 30 μM KBrO_3_, induced a less dramatic impact in cells than the initial treatment. Although also retreated cells showed an overall reduction in their gene expression activity (Figure 7B), this effect was smaller than the one caused by the first exposure (Figure 7C). Among the cellular processes that did not change or were upregulated during the second exposure but not the first, were some modules of gene regulation (especially mRNA splicing), protein catabolism and – interestingly – mitochondrial as well as DNA repair-related pathways. The DNA damage-associated genes included known repair enzymes, such as APEX1 (Figure 7D) but also the mitochondrial exonuclease MGME1 required for the degradation of damaged mtDNA (23).

**Figure 7.**
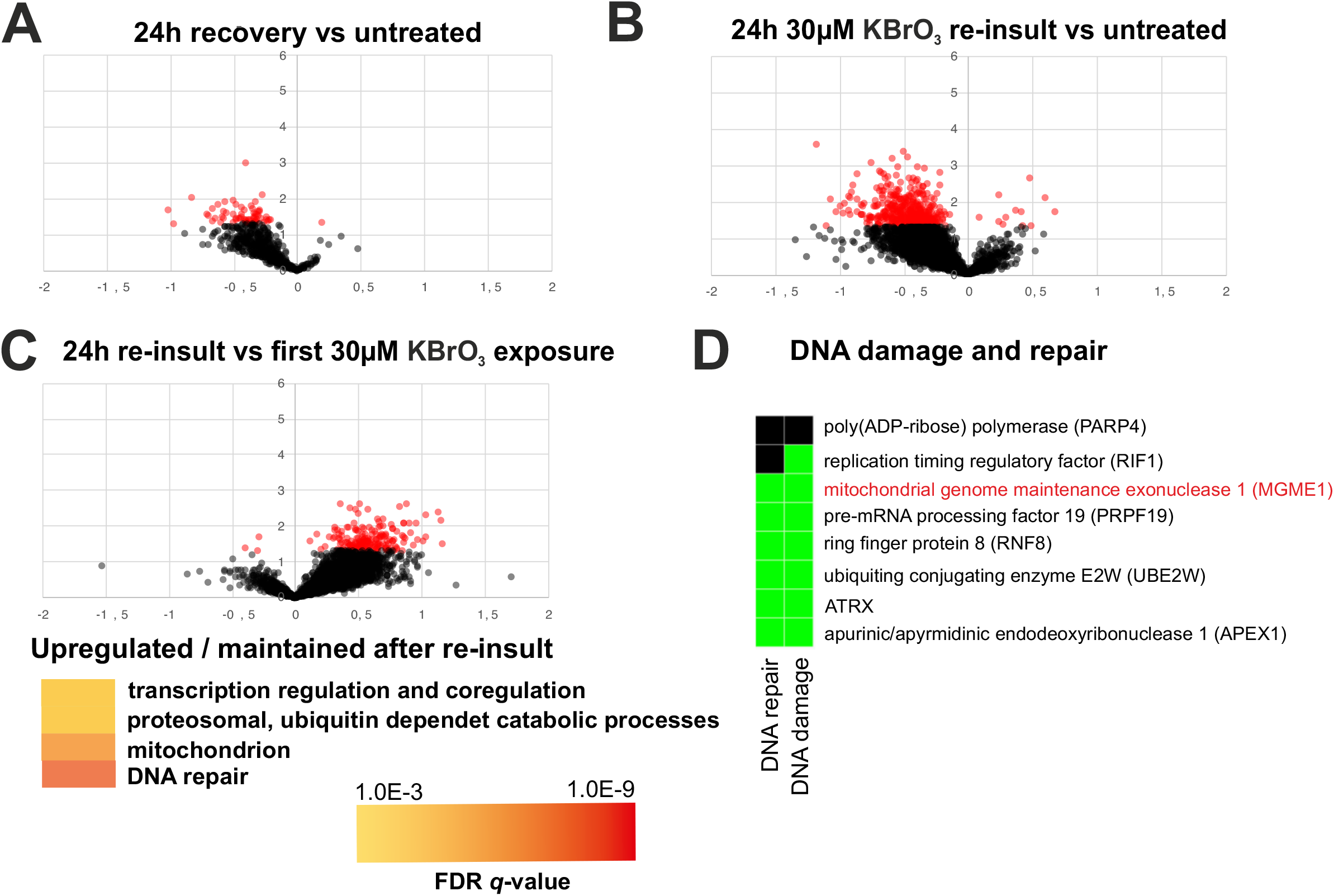
Hormetic effects in cells exposed to oxidative stress. (A) Gene expression differences between cells after 24h recovery of 30 μM KBrO_3_ treatment and untreated controls. The recovering cells still show several downregulated genes, but no specific pathways or processes are affected. (B) Retreatment with 30 μM KBrO_3_ after the recovery induces a similar, but milder reduction in gene expression as seen upon the first insult. (C) Retreated cells show the specific maintenance of pathways and cellular processes linked to gene expression, mitochondrial function and DNA repair. (D) Among the maintained DNA damage-associated genes are repair enzymes such as APEX1, but also the mtDNA-degrading exonuclease MGME1.

## Discussion

The study of free radicals and their effects in living organisms continues to be a hot topic in biology. Oxidative damage of biomolecules is detrimental and has been traditionally viewed as a pathological mechanism *per se* (3, 24). This is because pathological alterations of normal cellular functions, especially the dysfunction of mitochondrial electron transport chain typically results in increased oxidative stress, resulting in tissue damage. However, oxidative stress occurs frequently also under normal physiological conditions and plays an important regulatory as well as developmental role in most organisms. The balance between normal physiological regulation and pathology is highly interesting as it has evolutionary consequences on e.g. life history strategies (25). Most recently, there has been interest towards chronic oxidative stress caused by ionizing radiation during space exploration, which could have significance for the health of astronauts on prolonged spaceflights (26), including missions to Mars.

There are a number of ways to experimentally study oxidative stress. Most physiological might be the knockout or impairment of antioxidant defense genes (27), which also allows targeting oxidative stress to specific cell compartments or tissues (28). However, much of the work on generalizable, basic mechanisms is conducted using cultured cell models, where oxidative stress is induced by exposure to various chemicals. Some, such as the inhibitors of the ETC, are effective ROS generators, but cause major negative side-effects due to the poisoning of mitochondria (29). In this study, we have compared the effects of three oxidants that are commonly used in experimentation but are no direct ETC poisons and also have quite different modes of actions. While at the right concentrations all three, menadione, potassium bromate and hydrogen peroxide, are non-lethal but able to block cell proliferation (Figure 1C), there are subtle and more major differences in the oxidative stress they induce. One of the striking ones is the toxicity of menadione. We currently cannot explain why – as assumed – mitochondrial superoxide, originating from the bypass of electrons from ETC *via* menadione to oxygen, is able to cause a massive activation of nuclear DNA damage signaling, while KBrO_3_ or H_2_O_2_ are not (Figure 1C). Interestingly, menadione caused high levels of oxidative stress also in the cytoplasm, while the effect of KBrO_3_ is limited to mitochondria (Figure 1A). The mitochondrial ROS stress by KBrO_3_ correlates with its effects on mtDNA replication, while menadione does not influence mtDNA maintenance (Figure 6). This apparent mitochondrial specificity of KBrO_3_ is counterintuitive as the mitochondrial membrane potential and/or pH gradient should expel the negatively charged bromate from the organelle or retain it in the intermembrane space. However, the relatively small effects of H_2_O_2_ on cytoplasmic or mitochondrial ROS stress were not surprising due to the generally rapid turnover of the molecule (14).

One of the most important pathways for sensing ROS stress is the integrated stress response, or ISR (20). ISR was activated by all three oxidants, although the persistence and dynamics of the response differed (Figure 2). As with the activation of nuclear DNA damage signaling, menadione was the most aggressive of the tested chemicals, causing rapid and persistent activation of ISR already at low concentrations. Interestingly, ISR responded to KBrO_3_ with a delay, possibly indicating a slowly developing and chronic oxidative stress. Although we assumed that the ISR activation was PERK- and eIF2α-dependent, menadione-treated cells had steady-state levels of eIF2α phosphorylation, which were maintained also in the presence of a PERK inhibitor (Figure 5). Our observations underscore the notion that the textbook views of ISR and eIF2α phosphorylation probably do not represent the full picture, as pointed out recently (30). However, in line with the unfolded protein response-induced ISR (21), KBrO_3_-treated cells also showed a specific inhibition of protein synthesis apart for the transcriptional shutdown (Figure 4D).

Regardless of the nuances of ISR activation, menadione and KBrO_3_ treatments resulted in global downregulation of transcription (Figure 3B, C), as expected from a ROS-induced stress response (30). In contrast, the hydrogen peroxide treatment resulted in differential regulation of dozens of genes (Figure 3D). It is likely that our approach of treating the cells with one single large bolus of H_2_O_2_, causing an immediate damage and being subsequently lost in the various sinks, induces a different result than a persistent exposure to more stable concentrations of the oxidant, which could be achieved e.g. by a glucose oxidase approach (14). However, our experiment shows that the response to oxidative stress consists of more than just repair and recovery of the damage, which could reveal the mode of action of the different oxidants. While menadione-, KBrO_3_- and H_2_O_2_-exposed cells are all arrested at 24 h post treatment (Figure 1C, Figure 4B), only H_2_O_2_ cells are showing any adaptation or repair by amplifying their antioxidant defenses (examples in Figure 3D). Interestingly, these are mainly cytoplasmic enzymes, which together with the impacts on other cytosolic or extracellular metabolic functions (Figure 4F) suggests that extracellular H_2_O_2_ is unable to penetrate most cellular compartments. This also explains the lack of nuclear DNA damage signaling (Figure 1C). The fact that KBrO_3_-treated cells returned to the control state already 24 h after the removal of the drug (Figure 3A, 7A) also suggests that the effects seen in cells after 24 h H_2_O_2_ treatment are not only about recovery and repair. Despite the numerous H_2_O_2_ sinks in cells and medium, it is likely that the treatment sets in motion a cascade of events, interfering with a number of metabolic pathways, while not triggering a similar integrated stress response as menadione and potassium bromate. This observation emphasizes the fact that the differences in ROS responses are likely to be determined by a single or a set of specific oxidative reactions. Both the location and the order of oxidative reactions are plausible to make the difference between physiological signaling and an all-out stress response. It is worth pointing out that in this work we have focused on HEK293T cells, and that the ROS responses in other established cells lines, not to mention more natural primary cells or tissues, are likely to be different.

Finally, we looked at the potential hormetic effects induced by ROS exposure. We chose potassium bromate for this trial due to its lower toxicity compared to menadione, as it did not for example induce nuclear DNA damage response (Figure 1C). At the same time its dose was easier to control than H_2_O_2_, and it was able to cause stabile oxidative stress in cells, as judged from the ISR activation (Figure 2). The experiment should be considered as a proof-of-concept, as it is clear that mechanistic elucidation of the priming of hormesis would be a project on its own. However, our observations provide an interesting insight into the potential mechanisms of hormesis activation. Remarkably, although a second oxidative insult after recovery from an initial stress did induce silencing of gene expression (Figure 7B), corresponding to ISR activation, the response was muffled in comparison to the first exposure (Figure 3A, Figure 7C). While the re-treated cells did not upregulate any meaningful genes, they maintained the expression of groups of genes, which might enable adaptation to protein or DNA damage (Figure 7C). Among the DNA metabolism-linked gene products was the mitochondrial MGME1 nuclease, that is required for turnover of damaged mtDNA (23) and which we have recently shown to be associated with damage-induced changes in mtDNA replication (17). It is clear that the role of MGME1 in mtDNA damage response warrants further studies. The experiment also reveals interesting insight into the priming of the potential hormetic effects in cells. Notably, after recovery from the initial KBrO_3_ insult, the cells did not show significant alterations in gene expression patterns compared to the untreated (Figure 3A, 7A). While it is possible that the oxidative stress left behind some cellular memory in the form of protein modifications, a more plausible explanation are epigenetic modifications caused by the stress, which will influence the gene expression response in the following cell generations. Various types of stress, including oxidative stress (31), are known to induce epigenetic changes in cells. It is noteworthy that oxidative stress has been also linked with epigenetic changes seen in ageing individuals (32). While ageing is generally considered as deleterious to the individual, it is plausible that some of the epigenetic changes represent physiologically meaningful adaptations to oxidative metabolism and environmental stress.

As a conclusion, in the presented work we compared the effects of menadione KBrO_3_ and H_2_O_2_ on DNA damage response, integrated stress response and global gene regulation in HEK293T cells. Despite some similarities, the responses to the three oxidants differed markedly, urging caution when comparing studies using different sources of oxidative stress. We also observed that an initial oxidative stress can mitigate the consequences of a subsequent oxidative insult. Because the first oxidant exposure did not cause significant long-lasting effect on gene expression, it is plausible that the protection is caused by epigenetic modifications that have primed the cells to better tolerate or repair oxidative damage. The actual mechanisms of the hormetic effect should be investigated in future studies.

## Experimental procedures

### Cell culture and induction of oxidative stress

Human HEK293T cells were cultured in low glucose Dulbecco’s Modified Eagle Medium (DMEM, BioWest, L0103-500) with 10 % Fetal Bovine Serum (FBS) and 1 % Penicillin/Streptomycin. Cells were incubated at 37°C and 100% humidity with 8.5% CO_2_. To induce oxidative stress, cells were treated at 80 % confluency with menadione, H_2_O_2_ or KBrO_3_ using the concentrations and times given in the results. In experiments where a recovery phase was included, DMEM with the oxidizing agent was gently removed and replaced with fresh DMEM for up to 48 h.

For growth curve experiments, 10^5^ cells were seeded into 6-well plates and grown for 24 h before addition of the oxidizing agents. At 0 and 24 h, cells were detached with Trypsin/EDTA and counted using a Luna-FL cell counter (Logos Biosystems).

### roGFP constructs and measurements

The genes encoding cytosolic and mitochondrial matrix-targeted roGFP (33) were recloned from Addgene vectors 49435 and 49437 into pcDNA5FRT/TO and transfected into Flp-In T-REx 293 cells (ThermoFisher) to create stable cell lines with tet-inducible expression. These cells were grown in low glucose DMEM with 10% FBS, 100 ug/ml Hygromycin and 10 ug/ml blasticidin on 6 cm plates and roGFP expression was induced for 48h hours with 1 nM doxycycline before the addition of 100 μM H_2_O_2_, 10 μM Menadione or 30 μM KBrO_3_. After 4 h the medium was removed, the cells were detached with 1 ml PBS and 4x 200 ul were transferred to a black 96-well plate with clear bottom. GFP fluorescence was measured in a FluoStar Omega plate reader (BMG Labtech), using excitation wavelengths of 405 nm and 480 nm and an emission wavelength of >530 nm. Each treatment was measured in two biological replicates; non-induced cells served as blank, and induced, untreated cells as control. To verify the presence of equal cell numbers in the different conditions, the protein concentration of each well was determined by Bradford and a variability of <10% was found.

### MitoSOX measurements

Superoxide was quantified from live HEK293T cells using the fluorescent dye MitoSOX (ThermoFisher Scientific). 5 mM MitoSOX stock solution was prepared by dissolving one vial of MitoSOX (50 μg) in 13 μl DMSO. This stock was diluted further with 130 μl PBS. Cells treated with various oxidizing agents were washed with PBS, and 500 μl fresh medium with 5 μl MitoSOX was added, after which the cells were incubated 15 min at RT in the dark. Because HEK293T usually detached upon removal of DMEM/ MitoSOX and washing with PBS, the fluorescence was measured first in the MitoSOX-containing medium, then the cells were completely detached with medium, quickly spun down by centrifugation and resuspended in 200 μl PBS. The measurement was then repeated with this cell suspension. Fluorescence (absorption 400 nm/ emission 590 nm) was detected using a FLUOstar Omega microplate reader. To adjust for possible differences in cell density, the protein concentration of the resuspended samples was measured by Bradford assay and used as normalization factor.

### Protein extraction and Western blot analysis

Cells resuspended in 1 x PBS were spun down at 1000 x g at 4°C for 3 min. The PBS was removed, and the cell pellet was lysed in 4 × pellet volume of TotEx buffer (20 mM HEPES pH 7.9; 400 mM NaCl; 20% glycerol; 1% IGEPAL; 1 mM MgCl2; 0.5 mM EDTA; 0.1 mM EGTA; 10 mM β-Glycerophosphate; 10 mM NaF; 9 mM DTT and 1×complete EDTA-free protease inhibitor cocktail). 25-100 μg of the lysate was prepared for SDS-PAGE by mixing them with 14 volume 5 × Laemmli loading dye (250 mM Tris pH 6.8; 10 % SDS, 30 % glycerol, 0.5 M DTT; 0.02 % Bromophenol blue). Samples were denatured at 95°C for 5 min and separated over 12 or 15% Laemmli gels at 100 V for 90 min. The proteins were Western – blotted onto nitrocellulose membrane in Towbin buffer (25 mM Tris, 200 mM glycine; 0.1 % SDS, 20 % methanol) at 100 V for 90 min. The antibodies used in the study were: anti-VDAC (SigmaAldrich #SAB5201374), anti-Vinculin (SigmaAldrich #V9264), anti-ATF4 (Cell Signaling Technology #11815), anti-phospho-eIF2α (Ser51, Cell Signaling technology #3398) and anti-gH2AX (Biovision #3761).

### PERK inhibition

PERK was inhibited using 400 nM GSK2606414 (Merck Millipore). As of note, the used concentration is at the extreme high end of the tested concentrations for PERK inhibition (34), as we did not observe effects on eIF2α phosphorylation in HEK293T cells using lower concentrations of the inhibitor. For all the experiments, the inhibitor was added into the medium 4 h prior to the addition of the oxidative agent or UV exposure. The control UV exposure was performed as previously (16).

### mtDNA extraction and two-dimensional agarose gel electrophoresis (2D-AGE)

HEK293T cells were cultured in five 15 cm plates per condition. Cells were detached with medium and concentrated to 15 ml DMEM. 20 μg/ ml of Cytochalasin was added, and the cells were transferred to the incubator in a culture plate for 30 min incubation. The cells were transferred to a falcon tube and pelleted by centrifugation at 400 × g and 4 °C for 5 min. The pellet was resuspended in 5 ml H-buffer (225 mM mannitol, 75 mM sucrose, 10 mM EDTA, 10 mM HEPES-KOH pH 7,8, 1 mM DTT, 1 mg/ ml BSA) and cells were broken with 15 strokes in a tight dounce homogenizer. To remove nuclei and unbroken cells, the samples were centrifuged at 800 × g and 4 °C for 5 min, after which the centrifugation was repeated with the supernatant. Again, the supernatant was transferred, and mitochondria pelleted by centrifugation at 12,000 × g and 4 °C for 10 min. The pellet was resuspended in 1 ml H-buffer with BSA and DTT and the mitochondria purified using ultracentrifugation and two-step sucrose gradient (1/1.5 M sucrose on 20 mM HEPES pH 7.4 and 10 mM EDTA). The mitochondria were lysed in 1 ml mitochondrial lysis buffer (20 mM HEPES pH 7.4; 1 % SDS, 150 mM NaCl, 10 mM EDTA, 100 μg/ ml Proteinase K) for 20 min on ice. The lysate was then extracted with phenol-chloroform as described above, the DNA precipitated with 1 vol isopropanol and 20 μl 5M NaCl and the air-dried pellet was dissolved in 60 μl 20 mM HEPES pH 7.2.

For 2D-AGE (two-dimensional agarose gel electrophoresis), 5 μg mtDNA were digested with 3 ul *DraI* Fastdigest restriction enzyme (Thermo Scientific) at 37°C for 3h and extracted with 1 volume phenol-chloroform. The digested DNA was separated over a 0.4 % agarose gel in TBE at 55V overnight. To enable proper trimming for the second dimension, the 1D gel was stained with 1 μg/ml ethidium bromide in TBE and viewed under UV-light, so that the 1n fragment was visible. A 0.95% agarose second dimension in 1 × TBE with 1 μg/ ml ethidium bromide was cast around the cut 1D gel slices and separated overnight at 110 V with recirculation of TBE + 1 μg/ ml ethidium bromide at 4°C. After the run, the gel was inspected with UV light to confirm the success of agarose gel electrophoresis and blotted using standard procedures. The DNA fragment of interest was detected by prehybridizing the blot in Church’s buffer (0.25 M Na2HPO4 pH 7.4; 7 % SDS) at 65 °C for > 30 min, hybridized with a ^32^P-labelled human ND6 probe (nucleotides 14374-14595 of mtDNA) probe in Church’s buffer at 65 °C overnight and washed for 3x 15 min in 1x SSC, 0,1 % SDS. The radioactive signal was quantified with a Molecular Imager FX (BioRad) and Quantity One 4.6.2 software.

### RNA extraction, sequencing and transcriptome analysis

To study the effects of the different oxidants on the transcriptional regulation of the HEK293T cells, the cells were exposed either to 10 μM menadione, 30 μM potassium bromate or 100 mM hydrogen peroxide for 24 h. For recovery experiment the cells were exposed to 30 μM potassium bromate for 24 h, after which the medium was removed, cells washed once with fresh medium and let to incubate in refreshed medium for another 24 h. For re-treatment, the cells were split 1:3 after the first 24 h exposure to 30 μM KBrO_3_, let to recover in fresh medium for 24 h and treated again with 30 μM KBrO_3_ for another 24 h.

RNA was extracted from HEK293T using TRIzol (ThermoFisher Scientific), following the manufacturer’s instructions. RNA quality was assessed using the Agilent Bioanalyser, quantitated by Qubit (ThermoFisher) and high quality samples selected for analysis. RNA-Seq libraries were prepared using a multiplex 3’-capture method (35). Briefly, 10 ng of total RNA from each sample was tagged with an 8 base sample index and a 10 base unique molecular identifier (UMI) during initial poly(A) priming and reverse transcription. Samples were then pooled and amplified using a template switching oligonucleotide. The Illumina P5 (5’-AAT GAT ACG GCG ACC ACC GA −3’) and P7 (5’-CAA GCA GAA GAC GGC ATA CGA GAT −3’) sequences were added by PCR and Nextera transposase, respectively. The library was designed so that the forward read (R1) utilizes a custom primer (5’-GCC TGT CCG CGG AAG CAG TGG TAT CAA CGC AGA GTA C −3’) to sequence directly into the index and then the 10 base UMI. The reverse read (R2) uses the standard Illumina R2 primer to sequence the cDNA in the sense direction for transcript identification. Sequencing was performed on the NextSeq550 (Illumina), using the V2.5 High output kit generating 2 paired reads per cluster (19 bp R1; 72 bp R2). Adapters, primers, and low-quality bases were removed from the ends of raw reads using Trimmomatic v.0.34 (36). The resulting trimmed reads were mapped to the human genome (GRCh38) using STAR v.2.7.2(37) and the count table was created with the “`--quantMode GeneCounts`” option. The differential expression was performed using the gene count table in DESeq2 (38). Gene Ontology (GO) term enrichment analysis was performed using GOrilla (39) and DAVID Bioinformatics Resources 6.8 (40) online tools. The normalized transcriptome results are given in the supplementary material (**Supporting information file 1**).

## Data availability

All of data are contained within the manuscript and its supporting file.

## Supporting information

This article contains supporting information containing the transcriptome results (**Supporting information file 1**).

## Acknowledgements

MSc Maja Boziç is thanked for her assistance in the laboratory.

## Footnotes

## Abbreviations

2D-AGE: Two-dimensional Agarose Gel Electrophoresis
ETC: Electron Transport Chain
ISR: Integrated Stress Response
ROS: Reactive Oxygen Species
UPR: Unfolded Protein Response

